# TMEM120A contains a specific coenzyme A-binding site and might not mediate poking- or stretch-induced channel activities in cells

**DOI:** 10.1101/2021.06.17.448797

**Authors:** Yao Rong, Jinghui Jiang, Yiwei Gao, Jianli Guo, Danfeng Song, Wenhao Liu, Yan Zhao, Bailong Xiao, Zhenfeng Liu

**Author notes:** These authors contributed equally: Yao Rong, Jinghui Jiang and Jianli Guo. Corresponding author. (Zhenfeng Liu); (Bailong Xiao); (Yan Zhao).

## Abstract

TMEM120A, a member of the Transmembrane protein 120 (TMEM120) family, has pivotal function in adipocyte differentiation and metabolism, and may also contribute to sensing mechanical pain by functioning as an ion channel named TACAN. Here we report that expression of TMEM120A is not sufficient in mediating poking- or stretch-induced currents in cells, and have solved cryo-EM structures of human TMEM120A (*Hs*TMEM120A) in complex with an endogenous metabolic cofactor (coenzyme A, CoASH) and in the apo form. *Hs*TMEM120A forms a symmetrical homodimer with each monomer containing an amino-terminal coiled-coil motif followed by a transmembrane domain with six membrane-spanning helices. Within the transmembrane domain, a CoASH molecule is hosted in a deep cavity and forms specific interactions with nearby amino acid residues. Mutation of a central tryptophan residue involved in binding CoASH dramatically reduced the binding affinity of *Hs*TMEM120A with CoASH. *Hs*TMEM120A exhibits distinct conformations at the states with or without CoASH bound. Our results suggest that TMEM120A may have alternative functional roles potentially involved in CoASH transport, sensing or metabolism.

## Introduction

Through a fundamental cellular process known as mechanosensation, mechanical forces are converted into electrochemical signals during various physiological processes, such as touch, hearing, pain sensation, osmoregulation and cell volume regulation (García-Añoveros and Corey, 1997). In the past decades, various types of ion channels involved in sensing and transducing mechanical force signals have been identified (Jin et al., 2020). Among them, mechanosensitive channels of large and small conductances (MscL and MscS) serve as emergency release valves on the membrane by sensing and responding to membrane tension (Booth and Blount, 2012; Haswell et al., 2011). Piezo channels are involved in cellular mechanosensation processes crucial for touch perception, proprioception, red blood cell volume regulation and other physiological functions in animals (Murthy et al., 2017; Xiao, 2020). Besides, NompC from the Transient Receptor Potential (TRP) channel family, OSCA/TMEM63 and TMC1/2 from the TMEM16 superfamily, TREK/TRAAK channels from the two pore-domain potassium (K2P) channel subfamily and degenerins/acid-sensitive channels from the Epithelial Sodium Channel (ENaC) family also serve as mechanosensitive ion channels with pivotal functions in various mechanotransduction processes of eukaryotic organisms (Jin et al., 2020).

TMEM120 is a family of membrane proteins wide spread in animals and plants, and was originally identified as a nuclear envelope protein through a proteomic approach (Malik et al., 2010). Two paralogs of TMEM120 (TMEM120A and TMEM120B) exist in mammalian cells and they have crucial functional roles in adipocyte differentiation and metabolism (Batrakou et al., 2015). Knockout of *Tmem120a* in mice led to a lipodystrophy syndrome with insulin resistance and metabolic defects when the animals were exposed to a high-fat diet (Czapiewski et al., 2021). Recently, TMEM120A has been reported to function as an ion channel involved in sensing mechanical pain and thereby termed TACAN (Beaulieu-Laroche et al., 2020). The protein is expressed in a subset of nociceptors and associated with mechanically evoked currents in heterologous cell lines. Besides, TACAN protein exhibited nonselective cation channel activity when heterologously expressed in cell lines or reconstituted in liposome (Beaulieu-Laroche et al., 2020). Despite that TACAN is crucial for transduction of noxious mechanical stimuli, the mechanism of mechanosensation and ion permeation mediated by TACAN channel remains elusive. The molecular basis underlying the functions of TMEM120A in pain sensation and adipocyte differentiation are still unclear. Here we electrophysiologically characterized the ability of TMEM120A in mediating mechanically activated currents in cells and reconstituted proteoliposome membranes, and solved the cryo-electron microscopy (cryo-EM) structures of human TMEM120A (*Hs*TMEM120A) at two different conformational states. A striking discovery is revealed on the specific interactions between TMEM120A protein and an endogenous cofactor crucial for energy and fatty acid metabolism, namely CoASH (Sibon and Strauss, 2016), which might provide important insights into its bona fide functions in cells.

## Results

### Functional characterizations of TMEM120A

When expressed heterologously in cell lines including CHO, HEK293T and COS9 cells, *Hs*TMEM120A only appeared to cause pico-ampere level of increase in stretch-induced currents (Beaulieu-Laroche et al., 2020), prompting us to verify the role of TMEM120A in mediating mechanically activated (MA) currents. To exclude any endogenous Piezo1-mediated MA currents, we have chosen the previously reported Piezo1 knockout HEK293T cells (P1-KO-HEK) (Cahalan et al., 2015) as our heterologous expression system. Using a piezo-driven blunted glass pipette to mechanically probe the cell membrane under a whole-cell recording configuration, we recorded robust MA currents from the P1-KO-HEK cells transfected with either Piezo1 or Piezo2 (Fig. 1A, B). By contrast, similar to vector-transfected cells, none of the cells transfected with either mouse TMEM120A (*Mm*TMEM120A) or *Hs*TMEM120A showed poking-induced currents (Fig. 1A, B). These data suggest that TMEM120A is not sufficient to mediate poking-induced currents. To examine whether TMEM120A might specifically mediate stretch-induced currents as originally reported (Beaulieu-Laroche et al., 2020), we measured stretch-induced currents by applying negative pressure from 0 to −120 mmHg to the membrane patch under either cell-attached (Fig. 1C, D) or inside-out (Fig. 1E, F) patch-clamp configurations. As a positive control, Piezo1 mediated robust stretch-induced currents under both cell-attached and inside-out recording configurations (Fig. 1C, D). Consistent with the previous report that human TMEM63a (*Hs*TMEM63a) serves as a high-threshold mechanosensitive channel (Murthy et al., 2018), we indeed found that *Hs*TMEM63a-transfected cells reliably showed slowly activated stretch-induced currents, despite the maximal current amplitude is smaller than that of Piezo1-mediated currents (20.4±2.2 vs 64.1±7.4 pA) (Fig. 1C, D). By contrast, under either cell-attached or inside-out patch configurations, none of the 45 recorded TMEM120A-transfected cells showed stretch-induced currents, similar to vector-transfected cells (Fig. 1C, D). Thus, under our experimental conditions with the use of Piezo channels and *Hs*TMEM63a as proper positive controls, we conclude that TMEM120A is not sufficient to mediate poking or stretch-induced currents in the P1-KO-HEK cells.

**Figure 1.**
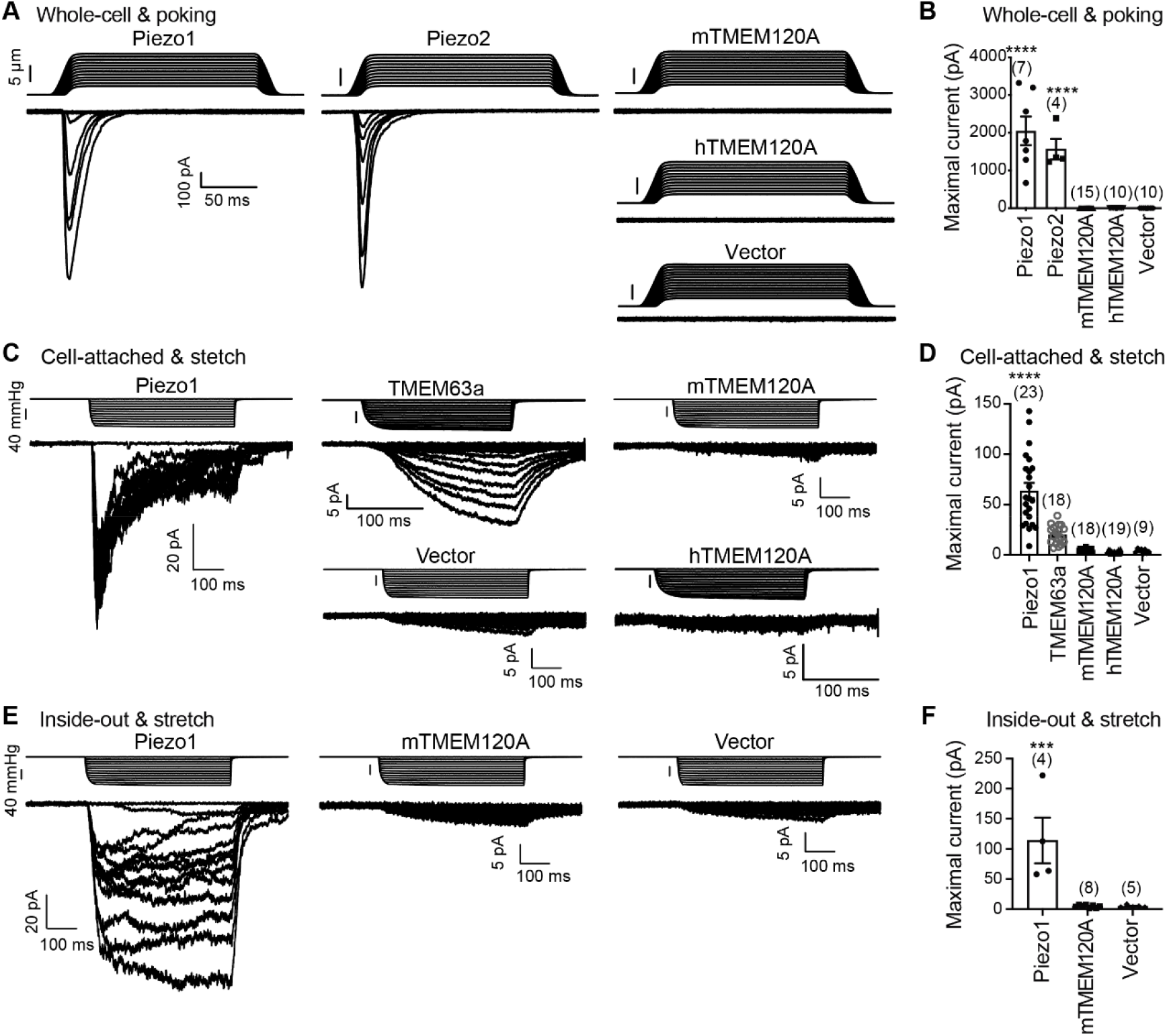
TMEM120A does not mediate poking- or stretch-induced currents in the P1-KO-HEK cells. **A**, Representative poking-evoked whole-cell currents from P1-KO-HEK cells transfected with the indicated constructs (*Mm*Piezo1-mRuby2, Piezo2-GST-ires-GFP, *Mm*TMEM120A-mCherry, *Hs*TMEM12A-mCherry, vector). **B**, Scatter plot of the maximal poking-evoked whole-cell currents. **C**, Representative stretch-induced currents from P1-KO-HEK cells transfected with the indicated constructs (*Mm*Piezo1-mRuby2, *Hs*TMEM63a-mCherry, *Mm*TMEM120A-ires-GFP, *Hs*TMEM120A-mCherry, vector) under the cell-attached patch configuration. **D**, Scatter plot of the maximal stretch-induced currents. **E**, Representative stretch-induced currents from P1-KO-HEK cells transfected with the indicated constructs (*Mm*Piezo1-mRuby2, *Mm*TMEM120A-ires-GFP, vector) under the inside-out patch configuration. **F**, Scatter plot of the maximal stretch-induced currents. In panels **B**, **D**, **F**, each bar represents mean ± s.e.m., and the recorded cell number is labeled above the bar. One way ANOVA with comparison to the vector. ***, P < 0.001; ****, P < 0.0001.

Purified TMEM120A proteins were showed to mediate spontaneous channel activities with an estimated single-channel conductance of ~250 pS, which is drastically different from the single-channel conductance of 11.5 pS measured in cells (Beaulieu-Laroche et al., 2020). To verify whether purified TMEM120A proteins might mediate channel activities when reconstituted into lipid bilayers (Beaulieu-Laroche et al., 2020), we have reconstituted purified *Hs*TMEM120A proteins into giant unilamellar vesicles (GUVs) and carried out single-channel recording on the inside-out excised membrane patch by applying negative pressure on the membrane. As shown in Fig. S1, out of over 200 patches measured, only two exhibited pressure-dependent mini-conductance channel activities. In one patch, the channel became more active in response to increasing negative pressure at both −80 and +80 mV (Fig. S1A), whereas the other patch only responded to increasing negative pressure at −80 mV but not at +80 mV (Fig. S1B). Most other patches were either silent and insensitive to the negative pressure applied (Fig. S1C), or exhibited leaky signals, suggesting that the overall channel activity of the protein reconstituted in GUV is fairly low.

Taken together, our functional characterizations did not verify the original report that TMEM120A might form a bona fide mechanically activated ion channel.

### Overall structure of TMEM120A

To further investigate whether TMEM120A might form an ion channel or not, we went on to solve the cryo-EM structure of *Hs*TMEM120A with the hope to provide hints about the function of TMEM120A through structural approach. As shown in Fig. 2 (and Fig. S2, Table S1), the cryo-EM structure of *Hs*TMEM120A protein reconstituted in lipid nanodiscs forms a homodimeric assembly with an overall shape resembling a seesaw rocker. The two monomers of *Hs*TMEM120A proteins are related by a two-fold symmetry axis running through their interface and perpendicular to the membrane plane. Each monomer contains an N-terminal soluble domain (NTD) on the cytosolic side and a transmembrane domain (TMD) embedded in lipid bilayer (Fig. 3A). There are two α-helices (intracellular helices 1 and 2, IH1 and IH2) in the NTD of each monomer (Fig. S3). The long α-helix (IH1) of NTD intertwines with the symmetry-related one and interacts closely with the shorter one (IH2). IH1, IH2 and their symmetry-related ones in the adjacent monomer collectively form a coiled-coil structure with the long axis running approximately parallel to the membrane plane (Fig. 3B). The TMD contains a bundle of six transmembrane helices (TM1-6) forming an asymmetric funnel with a wide opening on the intracellular side and a bottleneck on the extracellular side (Fig. 3D). Between NTD and TMD, there is a hinge-like motif (HM) with two long loops and a short amphipathic α-helix positioned near the membrane surface on the intracellular side. The HM intercalates at the gap between the TMDs of two adjacent monomers and serves to stabilize the dimer by interacting with both subunits (Fig. 3C). Evidently, the overall structure of *Hs*TMEM120A differs largely from the other mechanosensitive channels with known structures (Fig. S4), consistent with the previous analysis indicating it does not share sequence similarity with known ion channels (Arenas and Lumpkin, 2020).

**Figure 2.**
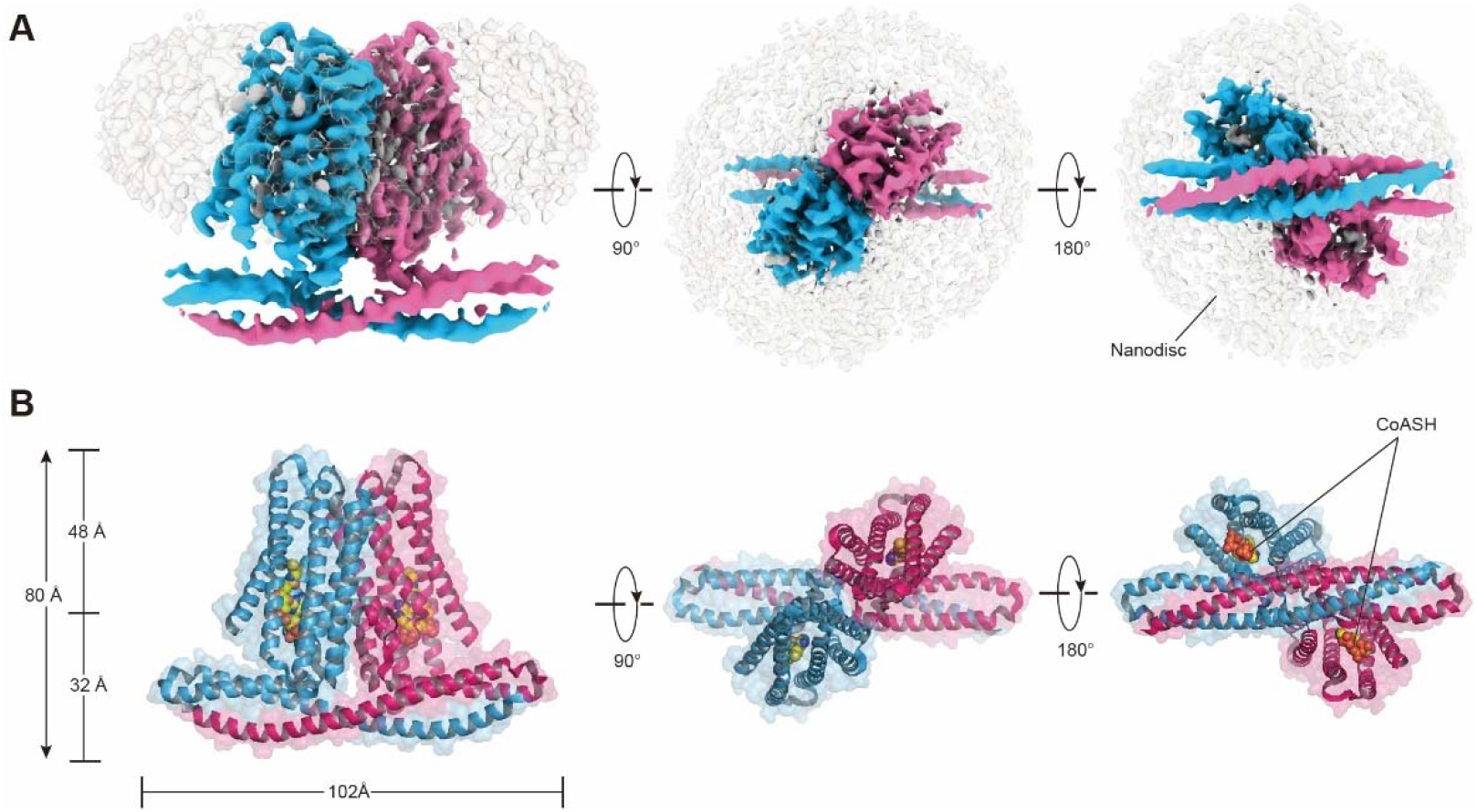
Cryo-EM density and overall architecture of *Hs*TMEM120A homodimer in complex with CoASH molecules. **A**, Cryo-EM density of *Hs*TMEM120A-CoASH complex dimer embedded in lipid nanodisc. The density of two *Hs*TMEM120A protein subunits are colored in blue and pink, while those of CoASH and lipid nanodisc are in silver. Side view along membrane plane, top view from extracellular side and bottom view from intracellular side are shown from left to right. **B**, Cartoon representations of the *Hs*TMEM120A-CoASH complex structure. The proteins are shown as cartoon models, whereas CoASH molecules are presented as sphere models. The views are the same as the corresponding ones in A.

**Figure 3.**
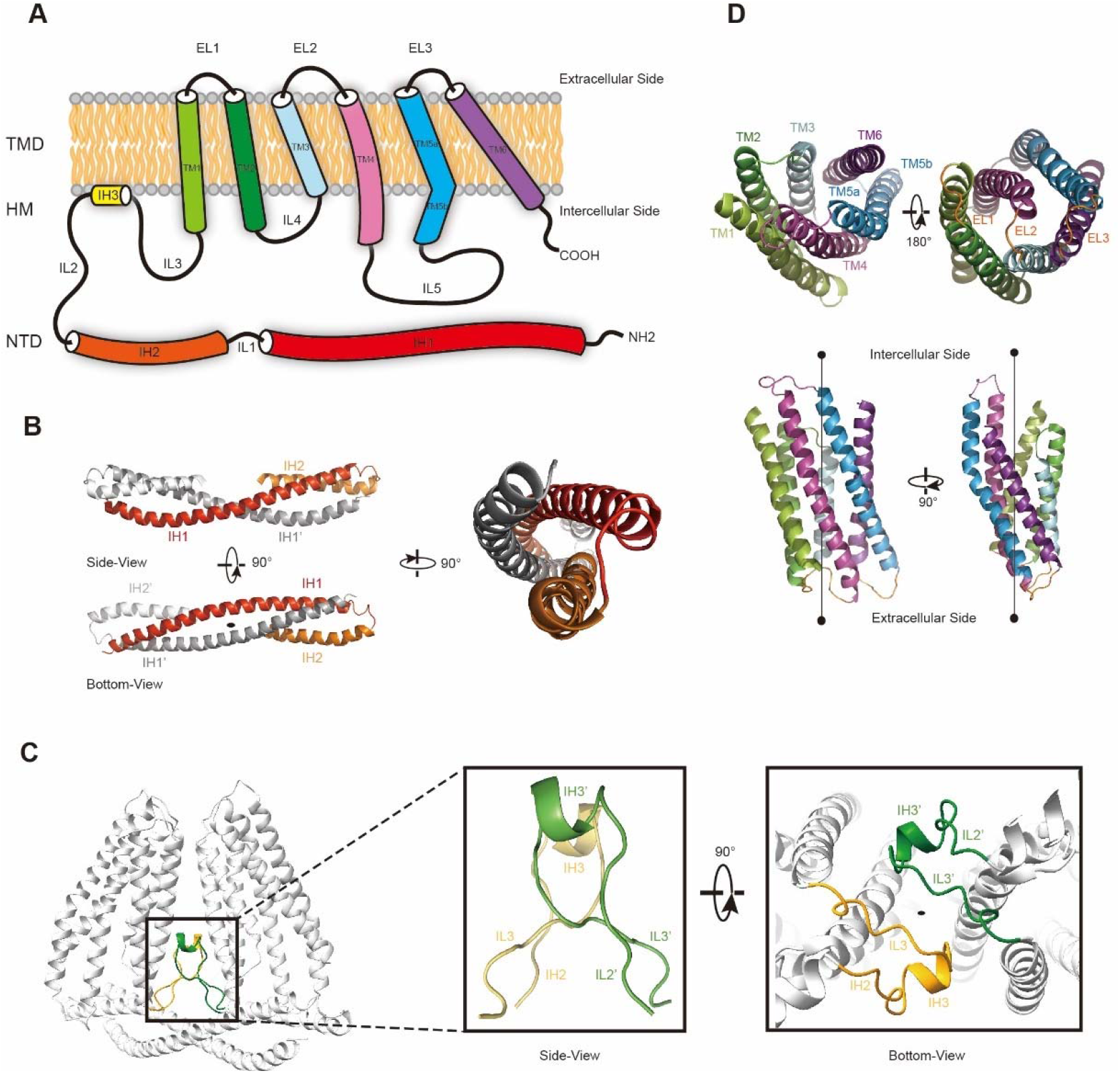
The membrane topology and domain structure of *Hs*TMEM120A monomer. **A,** The topology of *Hs*TMEM120A monomer and arrangement of different parts relative to the membrane. The α-helices are presented as cylinder models. IH1-3, the intracellular helices 1-3; TM1-6, transmembrane helices 1-6; IL1-5, intracellular loops 1-6; EL1-3, extracellular loops 1-3. **B**, The N-terminal domain (NTD) with two long and two short α-helices. **C**, The role of hinge-like motif (HM) in mediating dimerization of *Hs*TMEM120A at the monomer-monomer interface. The solid elliptical rings in B and C indicate the central two-fold axis of *Hs*TMEM120A homodimer. **D**, The transmembrane domain with a bundle of six transmembrane helices.

### Identification of a CoASH-binding site in HsTMEM120A

Unexpectedly, the *Hs*TMEM120A protein reconstituted in nanodiscs contains a CoASH molecule per monomer that is from endogenous cellular source and copurified along with the protein (Fig. 4A). Firstly, a small molecule density is observed in the intracellular funnel-like cavity of each monomer and the density matches well with the CoASH model, but not CoA derivatives (such as acetyl-CoA or fatty acyl CoA) or other nucleotide analogs (Fig. 4B). Secondly, mass spectrometry analysis on the perchloric acid extract of the purified *Hs*TMEM120A protein indicates that the protein does contain CoASH (Fig. 4C). Thirdly, The CoASH molecule can bind to the ligand-free *Hs*TMEM120A protein with a dissociation constant (Kd) at 0.685±0.045 μM (Fig. 4D). The endogenous CoASH in the protein sample can be removed by purifying the protein through the size-exclusion chromatography in detergent solution (see methods for details), as indicated by the decrease of A_260_/A_280_ (absorbance at 260 nm/absorbance at 280 nm) value from 0.88 to 0.58 (note: while the protein absorption maximum is at ~280 nm, the absorption maximum of CoASH is at ~260 nm (Zarzycki et al., 2008)). As shown in Fig. 4B, the CoASH molecule assumes a bent conformation and forms numerous specific interactions with amino acid residues on the lumen surface of the cavity (Fig. 4E). The amino acid residues involved in binding CoASH are highly conserved among various TMEM120A and TMEM120B homologs from different species (Fig. S5, S6). The 5-pyrophosphate and 3-phosphate groups of CoASH are positioned at the entrance of the cavity, while the adenosine and cysteine groups of CoASH inserted deep in the intracellular cavity pocket of *Hs*TMEM120A in a key-lock mode (Fig. 4B, E). Trp193 of *Hs*TMEM120A forms a strong π-π stacking interaction with the adenine group of CoASH. Mutation of the Trp193 to Ala dramatically reduced the affinity between CoASH and the protein (Fig. S7). Therefore, the TMD of *Hs*TMEM120A harbors a specific CoASH-binding site, potentially related to the function of TMEM120A protein.

**Figure 4.**
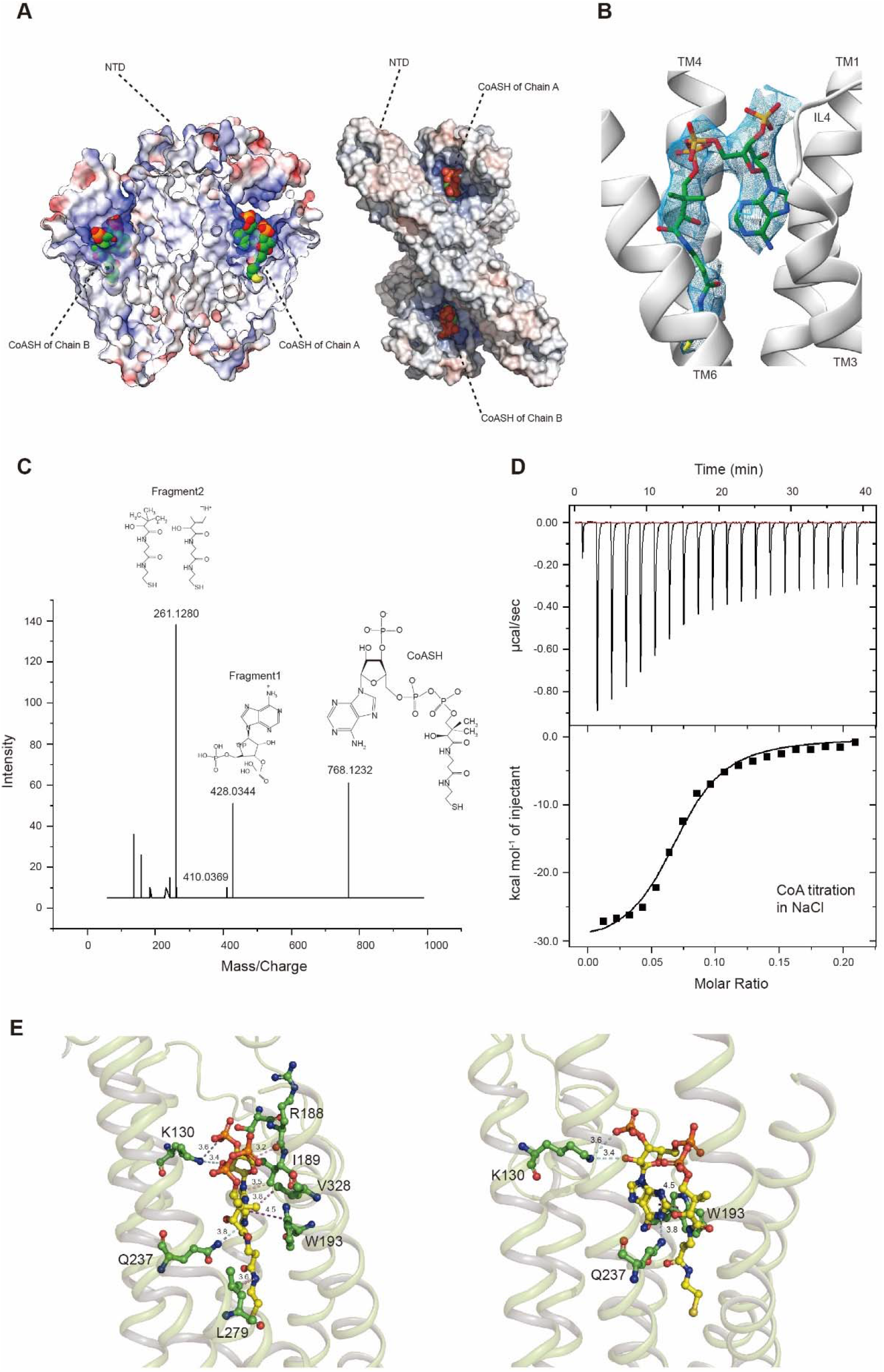
*Hs*TMEM120A contains an internal CoASH-binding site within each monomer. **A,** Electrostatic potential surface presentation of *Hs*TMEM120A dimer reveals a deep CoASH-binding cavity with electropositive surface. Left, side view; Right, top view along membrane normal from intracellular side. The CoASH molecules are presented as sphere models. **B**, The cryo-EM density of the ligand molecule bound to *Hs*TMEM120A fitted with a refined structural model of CoASH. **C**, Mass spectrometry analysis on the small molecule extracted from purified *Hs*TMEM120A protein. The chemical models of CoASH molecule and its fragements are shown above the corresponding peaks with m/z values of 768.1232, 428.0344 and 261.1280. **D**, Isothermal titration calorimetry analysis on the kinetic interactions between CoASH and *Hs*TMEM120A. The background heat of control was subtracted from the heat generated during binding of CoASH to *Hs*TMEM120A. The result is fit with the single-site binding isotherm model with Δ*H* = –31.09 ± 0.97 kcal/mol and *K*_d_ = 0.685 ± 0.045 μM. **E,** The detailed interactions between CoASH and the adjacent amino acid residues of *Hs*TMEM120A. Left and right, side views from two different angles. The blue dotted lines indicate the hydrogen bonds or salt bridge between Lys130 NZ and CoASH O2B (3.6 Å), Lys130 NZ and CoASH O8A (3.4 Å), Gln237 NE2 and CoASH N1A (3.8 Å). The purple dotted line shows the π-π interaction between Trp193 and CoASH at 4.5 Å distance. The pink dotted lines exhibit the non-polar interactions between CoASH and the adjacent residues (R188, I189, Leu279, and Val328) at <4 Å distance.

### Conformational change of *Hs*TMEM120A upon dissociation of CoASH

Does *Hs*TMEM120A change its conformation upon dissociation of CoASH? To address the question, we have solved the structure of *Hs*TMEM120A in detergent micelle with no CoASH bound (Fig. S8A-D, Table S1). The sample in detergent exhibits an A_260_/A_280_ ratio much lower than the one in nanodiscs (Fig. S8E, F), and no CoASH density is observed in the cavity (Fig. S8G, H). In the CoASH-free structure, the IL5 loop between TM4 and TM5 switches from the outward position to an inward position, partially covering the entrance of the intracellular cavity of *Hs*TMEM120A (Fig. 5A and B). The absence of CoASH creates a spacious vacant lumen in the intracellular cavity of *Hs*TMEM120A protein, potentially allowing water and ions to enter through the cytoplasmic entrance (Fig. S9A). Meanwhile, the cavity appears to be constricted at a narrow region on the extracellular side mainly by four amino acid residues, namely Met207, Trp210 and Phe219 and Cys310 (Fig. S9A). Among them, only Trp210 is highly conserved across different species, while the other three are less conserved (Fig. S5A and D, S6). In the presence of CoASH, the intracellular cavity of *Hs*TMEM120A is not only constricted by the four residues at the extracellular side, but also blocked by the bulky adenine group as well as the cysteine and pantothenate groups of CoASH (Fig. S9A and B).

**Figure 5.**
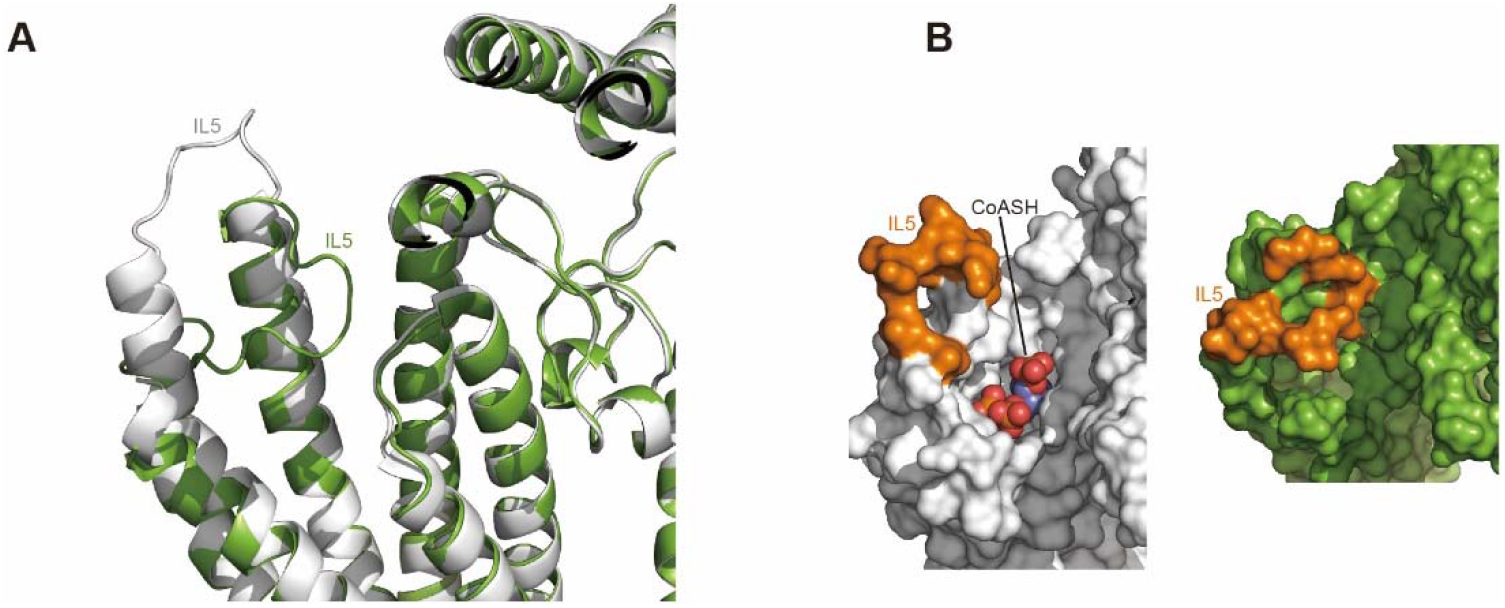
Structure of *Hs*TMEM120A at the CoASH-free state in comparison with the CoASH-bound state. **A,** Superposition of the structures of *Hs*TMEM120A at the CoASH-free state (green) and CoASH-bound state (silver). **B,** Surface presentation of the region around CoASH-binding site in the CoASH-bound (left) and CoASH-free (right) *Hs*TMEM120A structures. The IL5 loop region is highlighted in orange.

## Discussion

In the previous report (Beaulieu-Laroche et al., 2020), heterologously expressed TMEM120A only mediated picoampere-level of increase of stretch-induced currents on top of the endogenous currents, and failed to respond to poking stimulation.

However, it appeared to be more efficiently activated via a pillar-based mechanical simulation at the cell-substrate interface. The single-channel conductance of the TMEM120A-mediated currents measured in cells is 11.5 pS. Surprisingly, reconstituted TMEM120A proteins were showed to mediate spontaneous channel activities with an estimated single-channel conductance of ~250 pS (Beaulieu-Laroche et al., 2020). The drastically difference in the single-channel conductance measured in cells and reconstituted lipid membranes raises the concern whether TMEM120A itself might indeed form a bona-fide ion-conducting channel. Under our experimental conditions, when the purified *Hs*TMEM120A protein is reconstituted into GUVs, pressure-dependent channel activities were rarely observed during the single-channel recording processes. Moreover, we were unable to record TMEM120A-mediated poking or stretch-induced currents in cells (Fig. 1). Taken together, we conclude that TMEM120A is not sufficient to form an channel capable of sensing poking or stretch of the cell membrane. Nevertheless, we can not completely rule out the possibility that TMEM120A might only respond to certain forms of mechanical stimuli such as perturbation at the cell-substrate interface, which might require further independent verification. In case that TMEM120A does function as a mechanosensitive channel, the CoASH molecule may serve as a plug to block the intracellular entry and consequently stabilize the channel in a closed state, while the intracellular cavity may become accessible to ions upon dissociation of CoASH molecule from the cavity (Fig. S9C). When the channel is further activated by mechanical stimuli applied to the membrane, the constriction site on the extracellular side may open wider for ions to pass through the pore. However, the channel gating mechanism and coupling principle between putative CoASH inhibition and activation by mechanical stimuli during gating cycle would definitively need to be investigated further. Nevertheless, some other potential functions of TMEM120A, as a membrane-embedded enzyme utilizing CoASH as a substrate or cofactor, a CoASH transporter or a CoASH-sensing receptor, await to be explored further in the future.

## Methods

### Protein expression and purification

The cDNA encoding the full-length *Homo sapiens* TMEM120A (*Hs*TMEM120A) was synthesized with codon optimization (Genescript) for protein expression in the *Spodoptera frugiperda* Sf9 insect cells. The construct was cloned into pFastBac Dual vector between the 5’ EcoR1 and 3’ BamH1 sites, and an N-terminal FLAG followed by twin-Strep-tags is fused with the target protein. After purification, the plasmid DNA was used to transform DH10Bac cells to generate the recombinant bacmid. Positive clones containing desired genes were identified by blue/white selection and the recombinant bacmid were amplified and purified under sterile conditions. After three generations of amplification, the recombinant baculovirus were used for transfection of Sf9 cells adapted to suspension culture. To increase protein expression level, the Sf9 cells were cultured at 27 °C for 24 h and then at 20 °C for 48 h before being harvested.

The cells were harvested from the cell culture medium through centrifugation at 2,500 × g and disrupted by high pressure homogenizer (ATS Engineering Inc. Nano Homogenize Machine) under 1,200 bar for 6-8 cycles in a lysis buffer with 400 mM NaCl, 50 mM HEPES (pH 7.5), 10% glycerol and Protease Inhibitor Cocktail (MedChemExpress, Cat. No.: HY-K0010; 1 ml 100× stock solution per 150 ml solution). Unless stated otherwise, all steps of protein purification were conducted at 4 °C. The full-length *Hs*TMEM120A was extracted from cell membrane for 1 hour in a solubilization solution with 400 mM NaCl, 50 mM HEPES (pH 7.5), 1mM EDTA (pH 7.5), Protease Inhibitor Cocktail, 1% n-Dodecyl-β-D-Maltopyranoside (β-DDM) and 0.2% Cholesteryl Hemisuccinate (CHS). In order to remove insoluble fraction, the sample was centrifuged at 39,191 *g* (JL-25.50 rotor, Beckman) for 30 minutes.

The supernatant was incubated with the Streptactin beads for 2 hours and then the beads were loaded into a 20 ml chromatography column. The flow through was discarded and 5 column volumes (CV) of washing buffer with 50 mM NaCl, 20 mM HEPES (pH 7.5), 1 mM EDTA (pH 7.5), Protease Inhibitor Cocktail (MedChemExpress, 1 ml 100× stock solution per 150 ml solution), 0.006% glyco-diosgenin (GDN), 0.006% CHAPS and 0.001% CHS was applied to the column to remove contaminant proteins. Subsequently, the target recombinant protein was eluted in the elution buffer (5 CV) containing 50 mM NaCl, 20 mM HEPES (pH 7.5), 1 mM EDTA (pH 7.5), Protease Inhibitor Cocktail (MedChemExpress, 1 ml 100× stock solution per 150 ml solution), 0.006% GDN, 0.006% CHAPS, 0.001% CHS and 2.5 mM d-Desthiobiotin. The purified protein sample was concentrated in 100 kDa molecular weight cut-off (MWCO) concentrator (Millipore) to 3 mg/ml by centrifugation at 2,500 *g*.

The presence of CoASH in the purified *Hs*TMEM120A sample was monitored by measuring the absorbance under 260 nm and 280 nm (A_260_ and A_280_) of the protein sample with Nanodrop 2000 (Thermo). As the absorption peak of CoASH (copurified with the protein) is at ~260 nm(Zarzycki et al., 2008), relatively high A_260_/A_280_ value of 0.88-0.96 was measured with the purified *Hs*TMEM120A samples from different preparations. To further improve the homogeneity of purified protein, the concentrated protein sample was loaded onto a Superdex 200 increase 10/300 GL size-exclusion column (GE Healthcare Life Sciences) and eluted in the gel-filtration (GF) buffer containing 50 mM NaCl, 20 mM HEPES (pH 7.5), 1 mM EDTA (pH 7.5), Protease Inhibitor Cocktail, 0.006% GDN, 0.006% CHAPS and 0.001% CHS. The Peak fraction was collected and concentrated to 2.5-3.5 mg ml^−1^ in 100 kDa MWCO Amicon concentrator (Millipore). The A_260_/A_280_ value of the protein is lowered to 0.58-0.62 after size-exclusion chromatography.

For reconstitution of the *Hs*TMEM120A protein in nanodiscs, the protocol of protein solubilization and binding to the Streptactin resin as well as column washing was the same as the one described above. During the washing step, 5 ml washing buffer was added into the column so that the final volume of the mixture makes ~10 ml (containing 5 ml Streptactin beads). For nanodisc preparation, membrane scaffold proteins 1E3D1 (MSP1E3D1) was expressed in *Escherichia coli* cell, purified through immobilized metal affinity chromatography, and the His-tag was cleaved by the Tobacco Etch Virus (TEV) protease. The optimal ratio of *Hs*TMEM120A monomer : MSP1E3D1 : phospholipid is 1 : 2 : 150 (molar ratio). Initially, 2 ml of 10 mM phospholipid mixture (DOPE : POPS : POPC = 2 : 1 : 1, molar ratio, solubilized in 1% GDN and 2% β-DDM with ddH_2_O) was added into the column loaded with *Hs*TMEM120A protein and incubated for 30 minutes with constant rotation. Subsequently, the scaffold protein (MSP1E3D1) and Bio-beads (100 mg) were added into the column to trigger formation of nanodiscs by removing detergents from the system. The mixture was incubated for 2 hours with constant rotation. Another batch of Bio-beads (50 mg) was subsequently added and the mixture was incubate for 1 more hour. To remove the residual components and empty nanodiscs (without target protein), the column was mounted vertically on the stand and the solution on the column was separated from the resin through gravity flow. Afterwards, the resin on the column was washed with 10 CV of washing buffer with 50 mM NaCl, 20 mM HEPES (pH 7.5), 1 mM EDTA (pH 7.5) and Protease Inhibitor Cocktail. The *Hs*TMEM120A-nanodisc complex assembled on the Streptactin beads was eluted by applying 5 CV of elution buffer with 50 mM NaCl, 20 mM HEPES (pH 7.5), 1 mM EDTA (pH 7.5) and Protease Inhibitor Cocktail. The purified nanodisc sample was concentrated with 100 kD MWCO concentrator to 3.5 mg/ml through centrifugation at 2,500 *g*. To analyze the homogeneity of nanodiscs, the sample was run through size-exclusion chromatography (Superdex 200 increase 10/300 GL) and eluted in a solution with 50 mM NaCl, 20 mM HEPES (pH 7.5), 1 mM EDTA (pH 7.5) and Protease Inhibitor Cocktail. The major peak fraction was collected and concentrated to 3.5-4.5 mg ml^−1^ in 100 kDa MWCO Amicon concentrator. The purity of the protein sample was assessed by SDS-PAGE.

### Cryo-EM sample preparation and data collection

For Cryo-EM sample preparation, 3 μl of purified *Hs*TMEM120A sample was applied to the grid glow-discharged in H_2_/O_2_ for 60 seconds (Gatan Solarus 950). The Quantifoil 1.2/1.3-μm holey carbon grids (300 mesh, copper) were selected for preparing the *Hs*TMEM120A sample in detergent and 200 mesh grids of the same type were used for *Hs*TMEM120A-nanodisc sample. After being applied to the grids, the samples were blotted for 8 seconds with a Vitrobot (Vitrobot Mark IV, Thermo Fisher) after waiting for 10 seconds in the chamber at 4 °C with 100% humidity. The grid was plunge frozen in liquid ethane pre-cooled by liquid nitrogen. Cryo-EM images of the *Hs*TMEM120A particles in detergent were collected on a Titan Krios transmission electron microscope operated at 300 kV. The images were recorded by using SerialEM program(Mastronarde) at a nominal magnification of SA29000 in super-resolution counting mode and yielding a physical pixel size of 0.82 Å (0.41-Å super-resolution pixel size) using a K3 Summit direct electron detector camera (Gatan). The total dose on the camera was set to be 60 e^−^ Å^−2^ on the specimen and each image was fractioned into 32 subframes which were recorded with a defocus value at a range of −1.2 μm to −1.8 μm. The nominal magnification for the images of *Hs*TMEM120A-nanodisc sample was set to be SA22500 in super-resolution counting mode and yielding a physical pixel size of 1.07 Å (0.535-Å super-resolution pixel size). All the images were recorded by using a high-throughput beam-image shift data collection methods(Wu et al., 2019).

### Image processing

Two datasets (Datasets 1 and 2) were collected for the *Hs*TMEM120A nanodisc sample. For Dataset 1, a total of 4169 cryo-EM movies were aligned with dose-weighting using MotionCor2 program (Zheng et al., 2017). Micrograph contrast transfer function (CTF) estimations were performed by using CTFFIND 4.1.10 program (Rohou and Grigorieff, 2015). The procedures described below were performed by using cryoSPARC v3.1 program (Punjani et al., 2017) unless stated otherwise. After manual inspection of the micrographs, 3940 were selected and 230 particles were picked manually from the micrograph and sorted into 2D classes. The best classes were selected and used as references for autopicking procedure. After the process, 3,837,005 particles were auto-picked and extracted using a box size of 256 pixels. Six rounds of 2D classification were performed to remove ice spots, contaminants, aggregates and obscure classes, yielding 269,838 particles for further refinement. Non-uniform (NU) refinement was performed with C2 symmetry imposed and resulted in a 6.2 Å cryo-EM density map.

In order to improve the map resolution, a sequential heterogeneous refinement approach was applied according to the protocol described previously (Zhang et al., 2017). In details, after the NU refinement, well-resolved and biased reference models were generated through ab-initio reconstruction. Heterogeneous refinement with well-resolved and biased references was carried out to obtain a better density map. Next, the improved density map was subjected to low-pass processing at 15 Å and 30 Å to generate low resolution models. They served as the references of resolution gradient for another heterogeneous refinement. Finally, the empty micelles map was generated in Chimera by using the volume eraser tool. To obtain a noise downscaled map, the whole protein-nanodiscs complex map was subtracted with the region of peripheral scaffold protein-lipid shell downscaled with a scale factor of 0.5. By using the noise downscaled map and empty scaffold-lipid shell as references for the next step of heterogeneous refinement, a 4.7 Å map was obtained from 29,085 particles. The above process is the first round of sequential heterogeneous refinement. To enlarge the dataset of good particles, the 269,838 particles produced by 2D classification were divided into 9 subgroups, so that the number of particles in each subgroup is similar to the good particles yielding the 4.7 Å map. The good particles were combined with each individual subgroups and every chimeric subgroup was further processed in a second round of sequential heterogeneous refinement. Subsequently, all good particles from each subgroup were combined for a third round of sequential heterogeneous refinement yielding a 4.2 Å density map from 93,224 particles. For Dataset 2, a similar procedure was applied to yield 389,789 good particles leading to reconstruction of a map at 3.8 Å. The two datasets of the *Hs*TMEM120A-nanodisc sample were merged and subject to an additional round of 2D classification, the final round of sequential heterogeneous refinement, CTF refinement and NU refinement with *C*2 symmetry applied. The combined data resulted in a 3.7 Å density map from 410,963 particles after the entire process.

For the *Hs*TMEM120A sample in detergent, a total of 12,156 movie stacks were collected, followed by motion correction using MotionCor2 (Zheng et al., 2017) and CTF estimation using GCTF (Zhang, 2016). Particles were picked using Gautomatch (Zhang, 2017) and Topaz (Bepler et al., 2019) programs. Multiple rounds of 2D and 3D classifications were performed to clean up particles. The first round of 3D classification generated 8 classes, the class 7 (21.5%) showed well-resolved structural features including transmembrane helices and intracellular amphipathic helices. A second round of multi-reference 3D classification was then performed to improve the quality of map, giving rise to a subset of 80.7% particles. Particles were subsequently imported to cryoSPARC (Punjani et al., 2017) and the Non-uniform Refinement was carried out to yield a map at 4.3 Å resolution. The final map was generated through a subsequent local refinement in cisTEM (Grant et al., 2018) and was reported at 4.0 Å resolution according to GSFSC criterion. The local resolution of the map was estimated using the MonoRes program integrated in the Scipion framework (Vilas et al., 2018).

### Model building, refinement and validation

*Ab initio* model building of *Hs*TMEM120A was carried out in COOT 0.8.9 (Emsley et al., 2010) by referring to the 3.7 Å cryo-EM map and the secondary structure prediction by PSIPRED 4.0 (Buchan and Jones, 2019). The cryo-EM densities for the transmembrane domain (TMD) was well defined and provide detailed features for model building and real-space refinement. Registration of amino acid residues was guided mainly by the clearly defined densities of residues with bulky side chains such as Trp, Phe, Tyr and Arg. The density of IL5 (residues 250-263) was insufficient for assignment of the side chains, only the backbone of the polypeptide chain was built for this region. For the N-terminal domain (residues 7-100), some local regions were not well resolved in the cryo-EM map, involving about 40% amino acid residues of the domain. While the backbones of these residues were built in the model by referring to the density feature at high contour level (0.6034 V or 5.13 rmsd), their side chains remain uninterpreted.

For the structure of *Hs*TMEM120A in detergent, the overall resolution of cryo-EM density map was lower than that of *Hs*TMEM120A-nanodisc complex. Nevertheless, the densities of transmembrane domain (TMD) and hinge motif (HM) are well-resolved, allowing assignment of ~60% amino acid side chains. The initial model of the transmembrane domain (residues 123-336) was built by referring to the corresponding domain of the *Hs*TMEM120A-nanodisc complex structure. Around 85 residues of the 213 residues in the transmembrane domain are tentatively built as alanine residues as there are no side-chain information in the map. As the density of the N-terminal domain shows almost only the polypeptide backbone without side chain information, the region (residues 7-100) was built as a poly-alanine model.

The models were refined against the corresponding cryo-EM maps by using the phenix.real_space_refine program (Adams et al., 2010) followed manual adjustment in COOT to improve the overall geometry and fitting of models with the cryo-EM maps. The final models of the two *Hs*TMEM120A structures contained the bulk region of the protein covering amino acid residues 7-336. The pore profiles of the two structures were calculated by using HOLE program (Smart et al., 1996) and the narrowest site is defined as the location of the smallest pore radius. The electrostatic potential surface representations were calculated by ChimeraX (Goddard et al., 2018; Pettersen et al.) or the APBS (Baker et al., 2001) plugin in PyMOL (The PyMOL Molecular Graphics System, Version 2.0 Schrödinger, LLC). The statistics for data collection and processing, refinement and validation of the two *Hs*TMEM120A structures are summarized in Table S1.

### Mass spectrometry

The endogenous ligand (CoASH) bound to *Hs*TMEM120A protein was extracted from the purified *Hs*TMEM120A protein sample by using perchloric acid (72%) as described in a previous study (Shurubor et al., 2017). The purified *Hs*TMEM120A protein (in detergent) with A_260_/A_280_ value of 0.88 was concentrated to ~2 mg/ml. To extract the ligand, 1ml *Hs*TMEM120A sample was mixed with perchloric acid to give the final concentration of perchloric acid at 5%. The mixture was incubated on ice for 10 minutes before being vortexed for 10-15 minutes and centrifuged at 21,000× *g* for 10 min at 4 °C. The supernatant was removed and used for mass spectrometry (MS) analysis immediately. The sample was analyzed by using a high-performance liquid chromatograph (Dionex Ultimate 3000) equipped with a high-resolution mass spectrometer (SCIEX TripleTOF 5600). The HPLC separation was carried out on a ACQUITY UPLC CSH C18 reversed phase column (2.1mm × 100 mm, 1.7 μm (Waters)) at a flow rate of 0.25 ml/min. The column was maintained at 30 °C. The mobile phase consists of two components, namely phase A (50 mM ammonium acetate aqueous) and phase B (ACN). All solvents are of LC/MS grade. The column was eluted for 0-3.1 min with a linear gradient from 0% to 5% B, 3.1-9.0 min from 5% to 25% B, 9.0-11.0 min from 25% to 40% B and 11.0-15.0 min held at 40% B, followed by an equilibration step from 15.1 to 22 min at 0% B. The MS analysis was performed in positive ion mode at a resolution of 3,000 for the full MS scan in an information-dependent acquisition (IDA) mode. The scan range for MS analysis was 50-1200 m/z with an accumulation time of 250 ms in the TOF MS type and 100 ms in product ion type. Ion spray voltage was set at 5.5 kV in positive mode, and the source temperature was at 600 °C. The ion source gas 1 and 2 were both set at 60 psi, and the curtain gas was at 35 psi. In the product ion mode, the collision energy was set at 35 V, while the collision energy spread was at 15 V. Analysis of the LC/MS data was performed by using Peakview 2.1 software. The identification of CoASH was achieved on the bases of three criteria, namely exact mass, product ion peaks pattern and isotope spectrum. Values for m/z were matched within 5 ppm to identify the CoASH. The exact mass is 768.1225 based on the protonated molecular ion [M^+^H]^+^, [C_21_H_36_N_7_O_16_P_3_S^+^H]^+^. The spectral data obtained from the MS/MS showed two main product ion peaks at m/z 428.03 and m/z 261.13, corresponding to the fragments of CoASH.

### Isothermal titration calorimetry

The *Hs*TMEM120A protein was purified in the detergent conditions as described above. After gel filtration, the concentrated protein at 4.6 mg/ml was dialyzed overnight at 4 °C against 2 L dialysis buffer containing 50mM NaCl, 20mM HEPES (pH 7.5), 1mM EDTA (pH 7.5), Protease Inhibitor Cocktail (MedChemExpress, 1 ml 100× stock solution per 150 ml solution), 0.006% GDN, 0.006% CHAPS and 0.001% Cholesteryl Hemisuccinate (CHS). After concentration, the molar concentration of *Hs*TMEM120A monomer is at 120 μM and A_260_/A_280_ is at 0.58, and the molar concentration of W193A mutant monomer is at 17.6 μM and A_260_/A_280_ is at 0.62. Such a low A_260_/A_280_ value is close to the value (~0.6) for pure protein, suggesting that the endogenous CoASH bound to *Hs*TMEM120A protein is largely removed during the purification process. For the isothermal titration calorimetry (ITC) assay, the powder of coenzyme A (sodium salt hydrate, Sigma-Aldrich) was dissolved in the dialysis buffer to a final concentration of 6 mM as the stock solution, and then subject to serial dilution for the specific experimental requirements. All solutions were filtered with 0.22 μm filter membrane and then degassed prior to use. Measurements of enthalpy change (ΔH°) upon CoASH binding were performed by a MicroCal ITC200 (Malvern). In order to ensure that the *Hs*TMEM120A protein was saturated with CoASH at a reasonable rate, the optimized ratio of protein to CoASH was adjusted to 1: 1 (mol: mol). The sample cell was rinsed repeatedly by the dialysis buffer and 350 μl protein sample was loaded carefully into the cell to avoid formation of any bubbles.

The injection syringe was filled with 120 μM CoASH solution for wild type *Hs*TMEM120A or 17.6 μM CoASH solution for W193A mutant. The experimental temperature was set at 20 °C. The total number of injections was 20 and the volume of each injection was 2 μl except that the first injection was 0.4 μl. The CoASH solution were titrated into the *Hs*TMEM120A protein solution and the mixture was stirred at 700 rpm. The experimental data were analyzed by MicroCal ITC200. The heat change of the last four injections was so weak that the protein was regarded as completely saturated with CoASH despite that the heat produced did not go to zero. Thus, the heat generated by the last injection was subtracted from the heat generated by each injection. As a control, the dialysis buffer was titrated with CoASH under identical experimental conditions and the weak background heat was also subtracted from the protein-CoASH binding data in the final results.

### Whole-cell electrophysiology and mechanical stimulation

The P1-KO-HEK293T cells (Cahalan et al., 2015) were grown in poly-D-lysine-coated coverslips in DMEM supplemented with 10% FBS, 100 units/ml penicillin and 10 μg/ml streptomycin at 37 °C and 5% CO_2_. DNA constructs including Piezo1-mRuby2, Piezo2-GST-ires-GFP, *Hs*TMEM63a-mCherry, *Mm*TMEM120a-mCherry, *Mm*TMEM120a-ires-GFP, hTMEM120a-mCherry and vector were transfected using Lipofectamine 2000 (Thermo Fisher Technology). 24 h or 36 h after transfection and prior to electrophysiology, the cells were briefly digested with trypsin and sparsely re-plated onto poly-D-lysine-coated coverslips to obtain individual cells for recording. Electrophysiological recordings normally started about 2 h later after re-plating the cells. The patch-clamp experiments were carried out with a HEKA EPC10 as previously described (Zhao et al., 2016). For whole-cell patch clamp recordings, the recording electrodes had a resistance of 2–6 MΩ when filled with the internal solution composed of (in mM) 133 CsCl, 1 CaCl_2_, 1 MgCl_2_, 5 EGTA, 10 HEPES (pH 7.3 with CsOH), 4 MgATP and 0.4 Na2GTP. The extracellular solution was composed of (in mM) 133 NaCl, 3 KCl, 2.5 CaCl_2_, 1 MgCl_2_, 10 HEPES (pH 7.3 with NaOH) and 10 glucose. All experiments were performed at room temperature. The currents were sampled at 20 kHz, filtered at 2 kHz using Patchmaster software. Leak currents before mechanical stimulations were subtracted off-line from the current traces. Voltages were not corrected for a liquid junction potential (LJP).

Mechanical stimulation was delivered to the cell during the patch-clamp being recorded at an angle of 80° using a fire-polished glass pipette (tip diameter 3–4 μm) as described. Downward movement of the probe towards the cell was driven by a Clampex-controlled piezo-electric crystal micro-stage (E625 LVPZT Controller/Amplifier; Physik Instrument). The probe had a velocity of 1 μm/ms during the downward and upward motion, and the stimulus was maintained for 150 ms. A series of mechanical steps in 1 μm increments was applied every 10 s and currents were recorded at a holding potential of −60 mV.

### Cell-attached electrophysiology

Stretch-activated currents were recorded in the cell-attached or inside-out patch-clamp configuration using the HEKA EPC10 and the Patchmaster software as previously described (Zhao et al., 2016). The currents were sampled at 20 kHz and filtered at 1 kHz. The recording electrodes had a resistance of 2–5 MΩ when filled with a standard pipette solution consisting of (in mM) 130 NaCl, 5 KCl, 10 HEPES, 1 CaCl_2_, 1 MgCl_2_ and 10 TEA-Cl (pH 7.3, balanced with NaOH). The external solution used to zero the membrane potential consisted of (in mM) 140 KCl, 10 HEPES, 1 MgCl_2_, and 10 glucose (pH 7.3 with KOH). All experiments were carried out at room temperature. Membrane patches with a seal resistance of at least 2 GΩ were held at −80 mV and stimulated with 200 ms or 500 ms negative pressure pulses given at either 5 or 10 mmHg steps from 0 to −120 mmHg through the recording electrode using a Patchmaster-controlled pressure clamp HSPC-1 device (ALA-scientific).

### Single channel electrophysiology on proteoliposome vesicles

The purified *Hs*TMEM120A protein was reconstituted into the membrane of giant unilamella vesicles (GUVs) by using the mixture of cholesterol:DPhPC:azolectin (w:w:w) ratio of 1:5:17 or 1:10:15.) with protein:azolectin ratio at 1:85 (w/w) through a modified sucrose method(Battle et al., 2009). The bath and pipette solutions contained 500 mM NaCl, 10 mM CaCl_2_ and 10 mM HEPES-NaOH (pH 7.4). Patch pipettes with resistances of 5-6 MΩ were used, and the patch resistance increased to ~2 GΩ after the pipette sealed tightly with the GUV membrane. Negative pressure was applied through a Suction Control Pro pump (Nanion) with a stepwise or linear protocol while data were recorded at constant holding potential. The data were acquired at 50 kHz with a 0.5-kHz filter and a HumBug 50/60 Hz Noise Eliminator (Quest Scientific), using an EPC-10 amplifier (HEKA). The Clampfit Version 10.0 (Axon Instruments) was used for data analysis and Igor Pro 6.37A (WaveMetrics) was used for making the graphs.

## Acknowledgments

The cryo-EM data were collected at the Center for Biological Imaging (CBI), Core facilities for Protein Science at the Institute of Biophysics, Chinese Acadmy of Sciences. The sample screening time on Talos Arctica was sponsored by the National Laboratory of Biomacromolecules and CBI. We thank X.J. Huang, B.L. Zhu and X.J. Li at CBI for their assistance in cryo-EM data collection; Y.Y. Chen, Z.W. Yang and B.X. Zhou at IBP for their help with ITC experiments; N.L. Zhu and F.Q. Yang at the Laboratory of Proteomics, IBP for the mass spectrometry analysis on the sample extracted from purified *Hs*TMEM120A protein; X. B. Liang for her support in sample preparation and data collection. The project is funded by the National Natural Science Foundation of China (31925024 and 31670749 to Z.L.; 31825014 and 31630090 to B.X.) and the Strategic Priority Research Program of CAS (XDB37020101 to Z.L. and XDB37030304 to Y.Z.).

## Author Contributions

Y.R. designed the expression construct of *Hs*TMEM120A, expressed and purified the *Hs*TMEM120A protein and reconstituted it in nanodiscs, extracted and identified CoASH through biochemical approaches, prepared the cryo-EM grids, collected cryo-EM data, built the model and refined the structure and carried out the ITC experiments. J.G. reconstituted HsTMEM120A in GUV and performed single-channel recording experiments. D.S. processed the cryo-EM data of *Hs*TMEM120A-nanodisc sample; Y.G and Y.Z. processed the cryo-EM data of HsTMEM120A-detergent sample; J.J. and W.L. carried out electrophysiolocal recordings in cells and analyzed data; Y.R., Y.Z. and Z.L. analyzed the structure. Y.R., J.G., Y.Z., B.X. and Z.L. wrote the manuscript. Y.Z., B.X. and Z.L. conceived and coordinated the project.

