## Supplemental Figures S1-9 and Table S1 for "TMEM120A contains a specific coenzyme A-binding site and might not mediate poking- or stretch-induced channel activities in cells"

**Supplementary Figures S1-9 and Table S1**

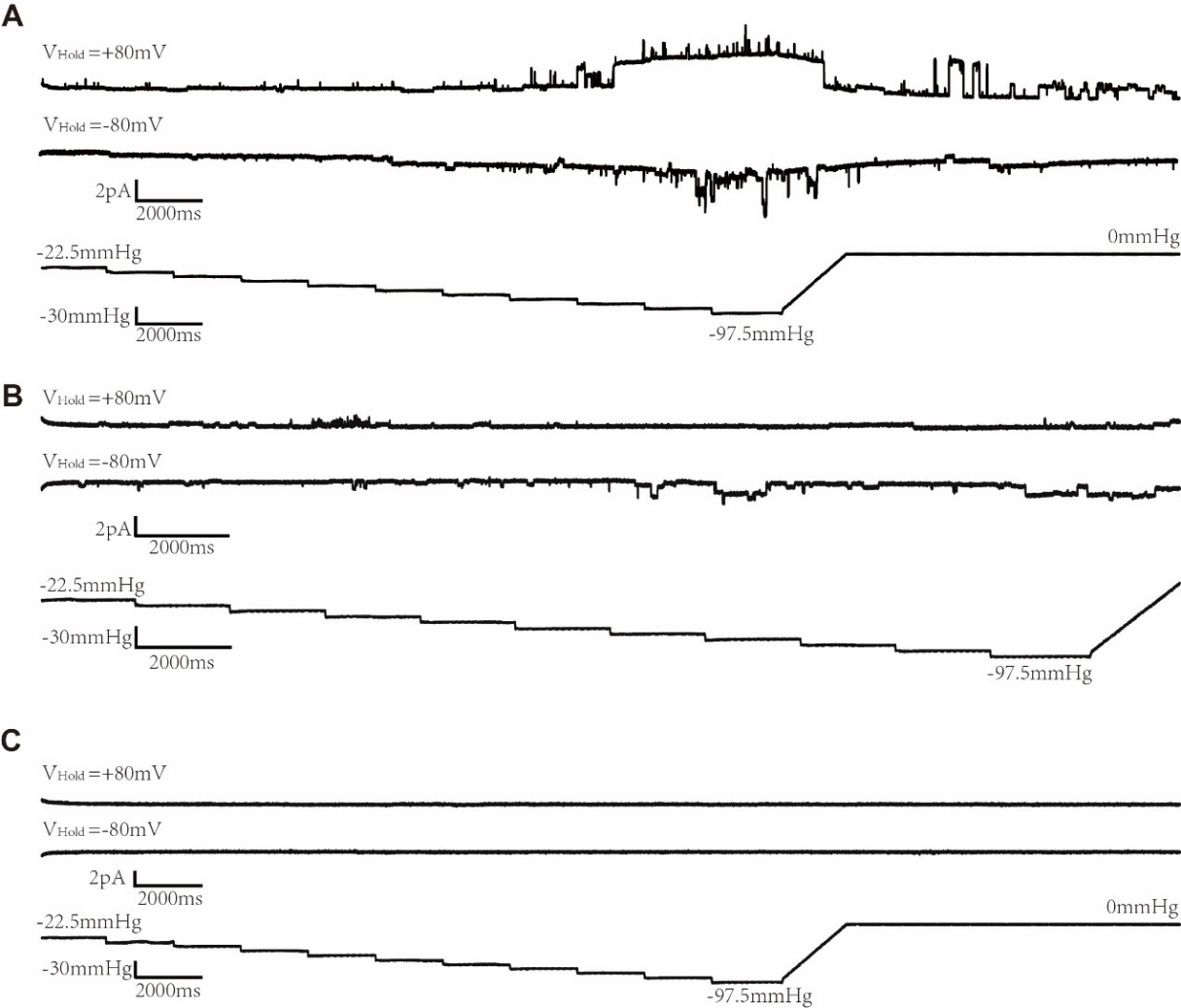

**Figure S1. Representative excised inside-out patch recordings of *Hs*TMEM120A reconstituted in GUVs.** **A,** Traces of ion channel activities responding to the increase of negative pressure at both +80 and -80 mV. **B,** A different patch of membrane exhibiting distinct behaviors of ion channel activities at +80 and -80 mV**.** While the trace recorded at -80 mV exhibits stimulated channel activities in response to increasing pressures, the one at +80 mV appears to be insensitive to increasing pressure. **C**, A silent patch with no pressure-dependent ion channel activities. The traces from three representative patches were obtained by holding the patch at -80 mV and +80 mV with a pressure pulse protocol shown at the bottom: -22.5 to -97.5 mmHg with -7.5 mmHg step. All recordings were performed with symmetrical pipette and bath solution containing 500 mM NaCl, 10 mM CaCl_2_ and 10 mM HEPES (pH 7.4 , ~300 Osm/L).

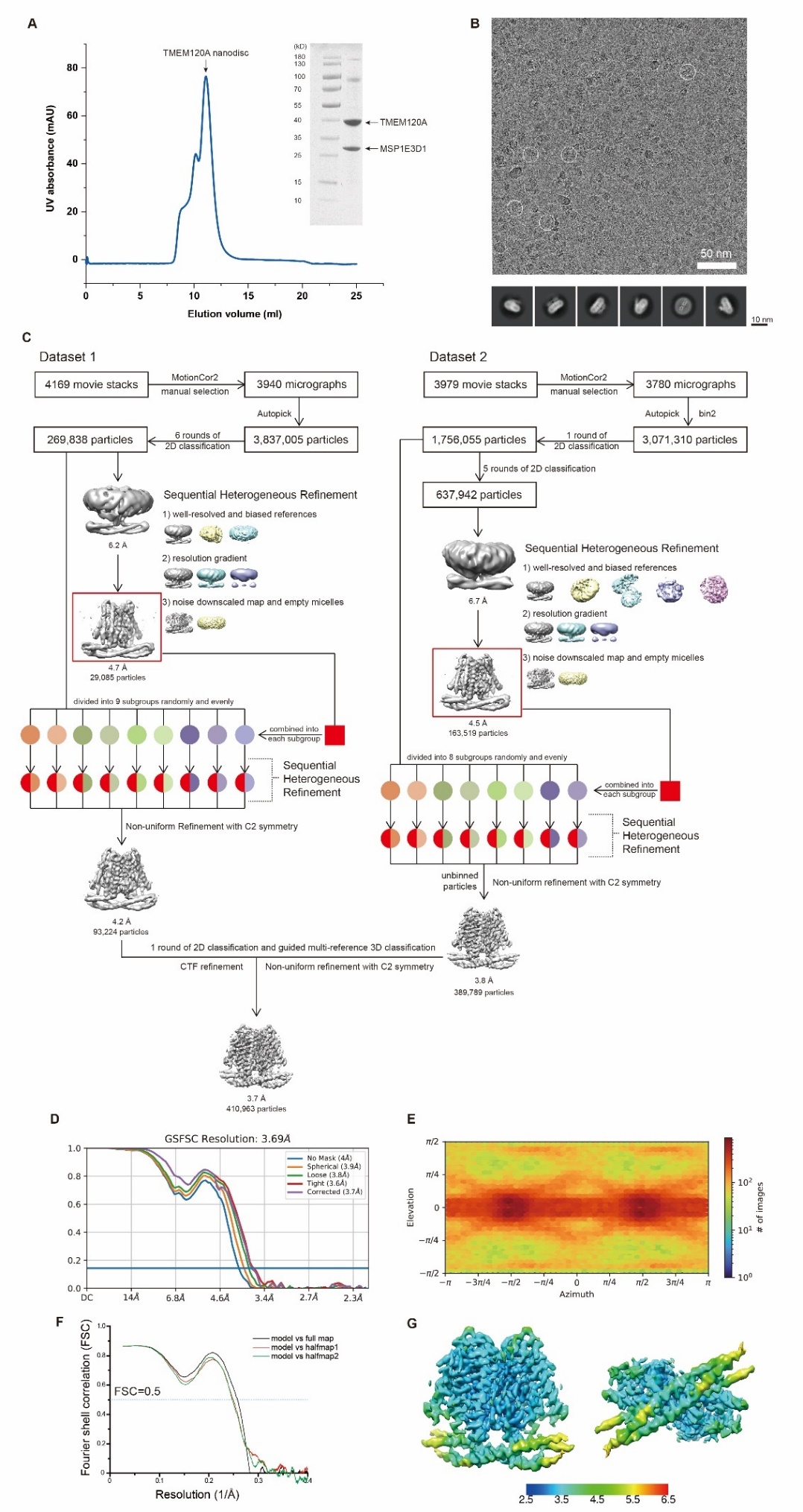

**Figure S2. Sample preparation, cryo-EM data collection and processing of the *Hs*TMEM120A-CoASH complex reconstituted in nanodiscs. A,** Size-exclusion chromatography of the *Hs*TMEM120A-CoASH complex reconstituted in nanodiscs. The image of SDS-PAGE analysis of the peak fraction is shown on the right. **B**, Representative cryo-EM image of the *Hs*TMEM120A-CoASH complex in nanodiscs and 2D classification images. **C,** The flow chart of data processing, 3D classification and refinement. **D,** The Gold Standard Fourier Shell Correlation (GSFSC) curves for the 3D refinement of the final cryo-EM map. Blue, The FSC curve obtained without application of mask; Orange, The FSC curve obtained with spherical mask; Green and red, The FSC curve obtained with loose and tight masks around the protein density respectively; Purple, The FSC curve obtained with corrected mask. **E**, Angular distribution of the particles used for the final reconstruction. **F**, The FSC curves between the structural model and full map as well as two half maps. **G,** Estimation of the local resolution of the final cryo-EM map. The unit for the numbers labeled nearby the gradient color bar is Å.

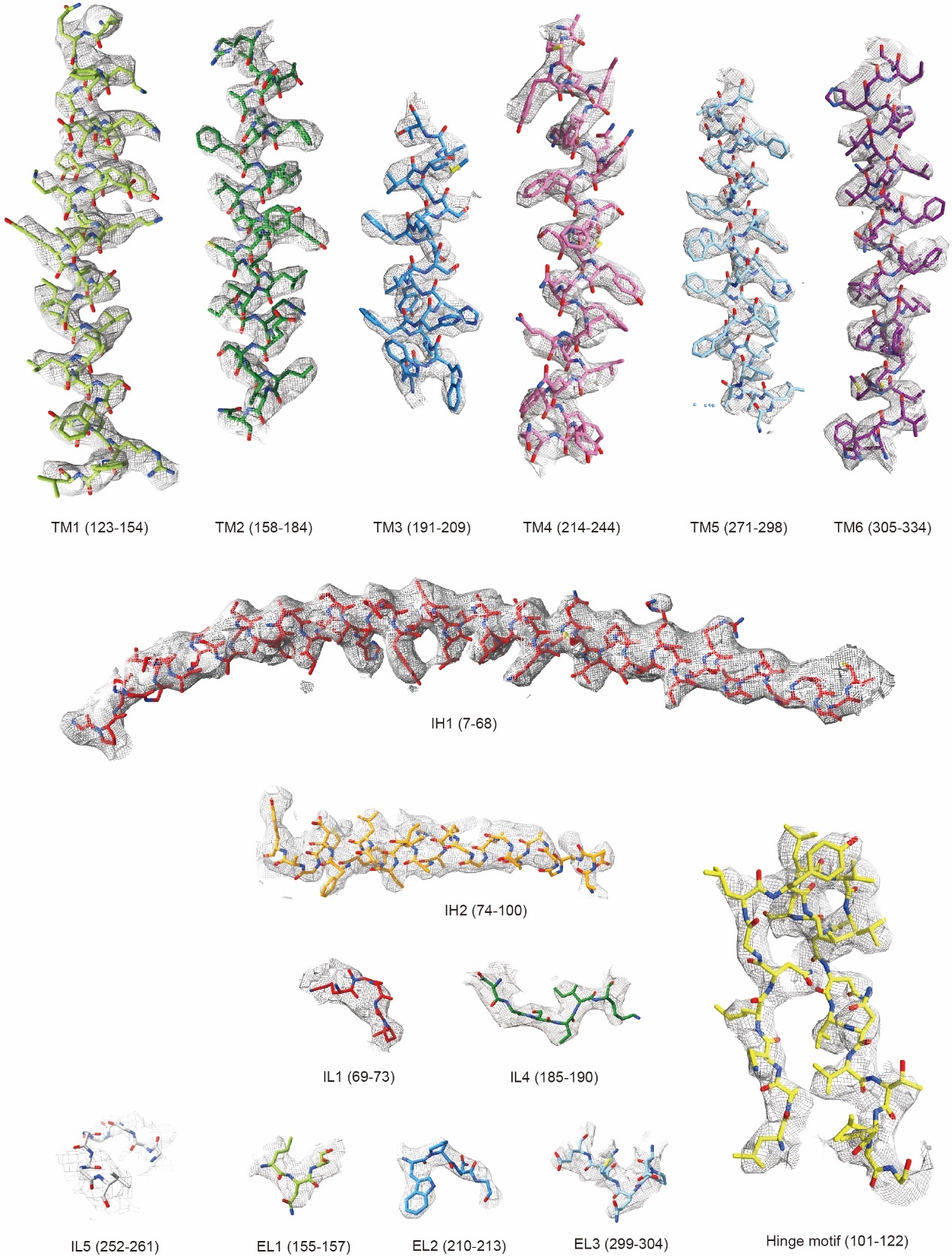

**Figure S3. Fitting of the structural model with the cryo-EM densities of various local regions of *Hs*TMEM120A in nanodiscs.** The refined structural model superposed on the map are shown as stick models. The map is contoured at 6.7-13.1 rmsd level. TM2 (158-184), TM3 (191-209), TM4 (214-244) and EL2 (210-213), 13.1 rmsd; hinge motif (101-122), TM1 (123-154) and TM5 (271-298), 11.0 rmsd; TM6 (305-334), EL1 (155-157) and IL4 (185-190), 9.1 rmsd; IH1 (7-68), IH2 (74-100), IL1 (69-73) and EL3 (299-304), 6.73 rmsd; IL5 (252-261), 4.33 rmsd.

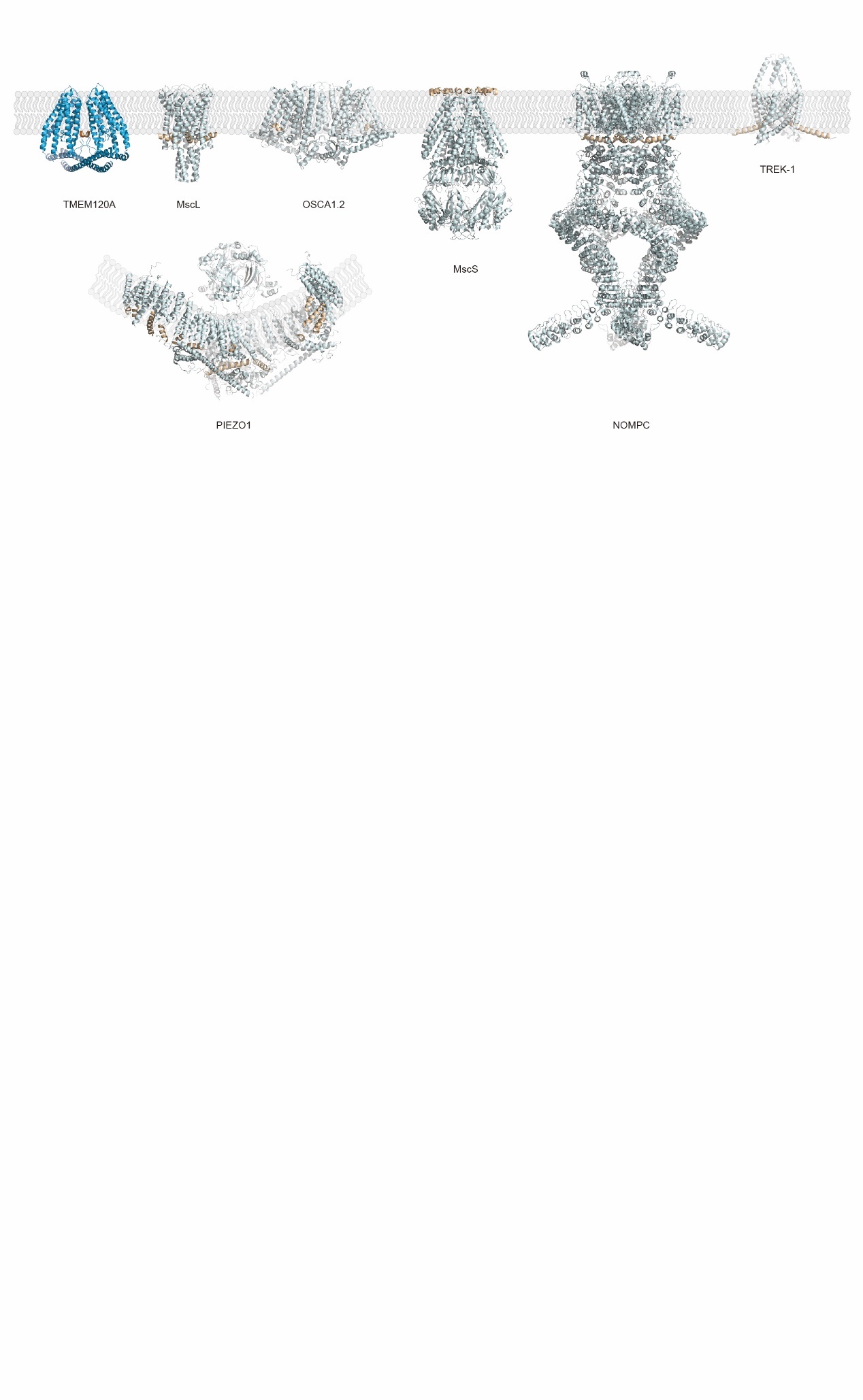

**Figure S4. Comparison of the *Hs*TMEM120A structure with those of various mechanosensitive channels.** MscL and MscS, mechanosensitive channels of large and small conductances (PDB codes: 2OAR and 6PWP); OSCA1.2, a plant hyperosmolality-gated Ca^2+^-permeable channel (PDB code: 6MGV); NOMPC, a drosophila mechanosensitive channel involved in gentle-touch sensation (PDB code: 5VKQ); Piezo, a mechanically activated channel mediating painful touch sensation (PDB code: 5Z10). TREK-1, a mechanosenstive two-pore domain K^+^ channel involved in polymodal pain perception, neuroprotection and general anesthesia (PDB code: 6CQ6).

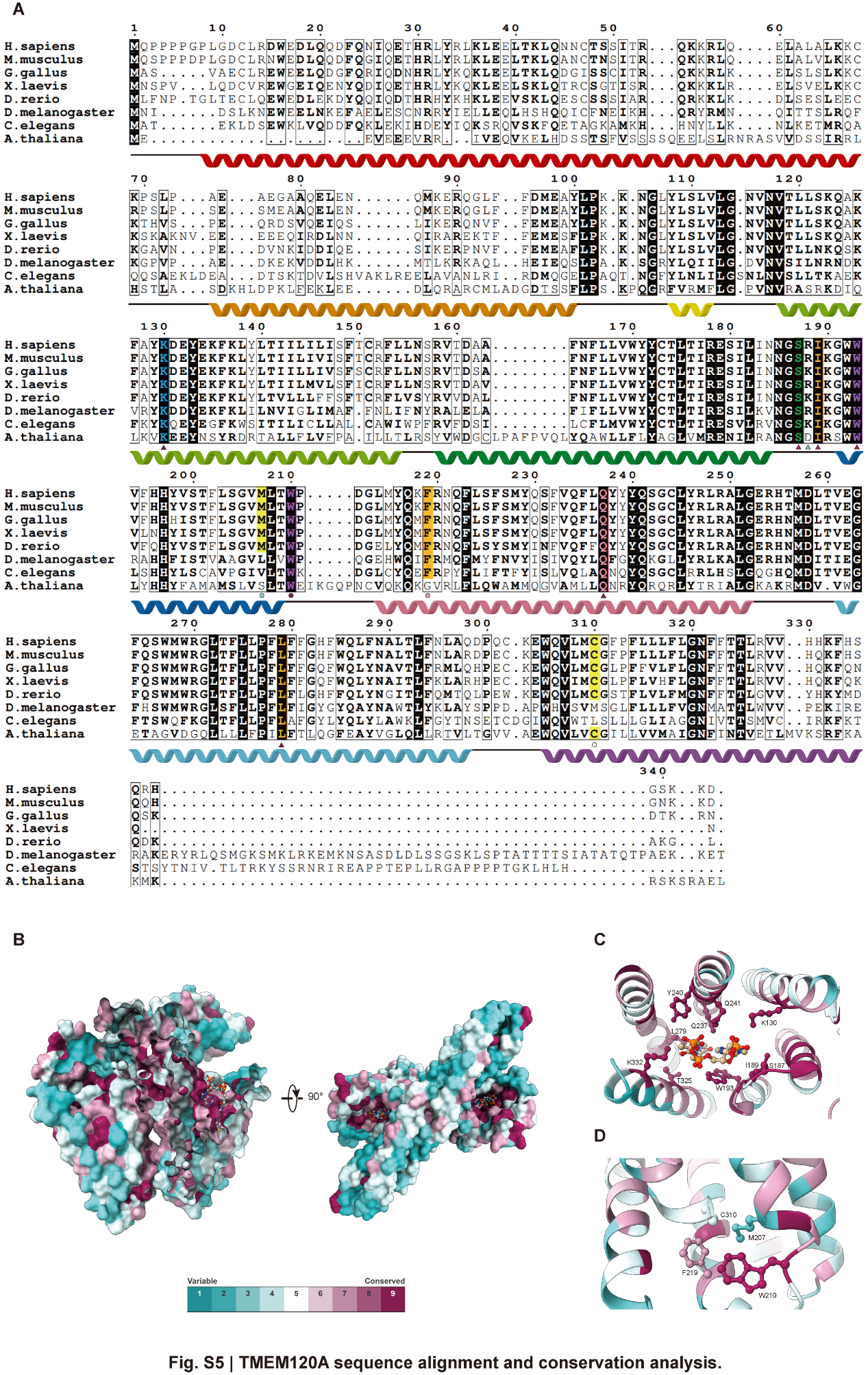

**Figure S5. The conserved features of TMEM120A. A,** Sequence alignment of *Hs*TMEM120A with other homologs from different species. The highly conserved amino acid residues are highlighted in dark background. The triangles denoted the CoASH binding residues and the color code is the same as the one for ConSurf conservation scores bar shown in B. The circles indicate the amino acid residues located in the narrowest region of channel pore. The α-helices and associated loops were indicated by the helical ribbons and dark lines labeled above the sequences respectively. Accession codes: *Homo sapiens* (NP_114131.1), *Mus musculus* (NP_766129.1), *Gallus gallus* (XP_040543083.1), *Xenopus laevis* (NP_001091170.1), *Danio rerio* (NP_001076452.1), *Drosophila melanogaster* (NP_001245509.1), *Caenorhabditis elegans* (NP_001370801.1) and *Arabidopsis thaliana* (CAD5314357.1). **B,** Mapping of the conserved amino acid residues on the surface of *Hs*TMEM120A structure. The conserved residues are shown in dark pink, while the variable ones are in cyan. **C** and **D**, The amino acid residues located at the CoASH-binding site and the constriction area respectively.

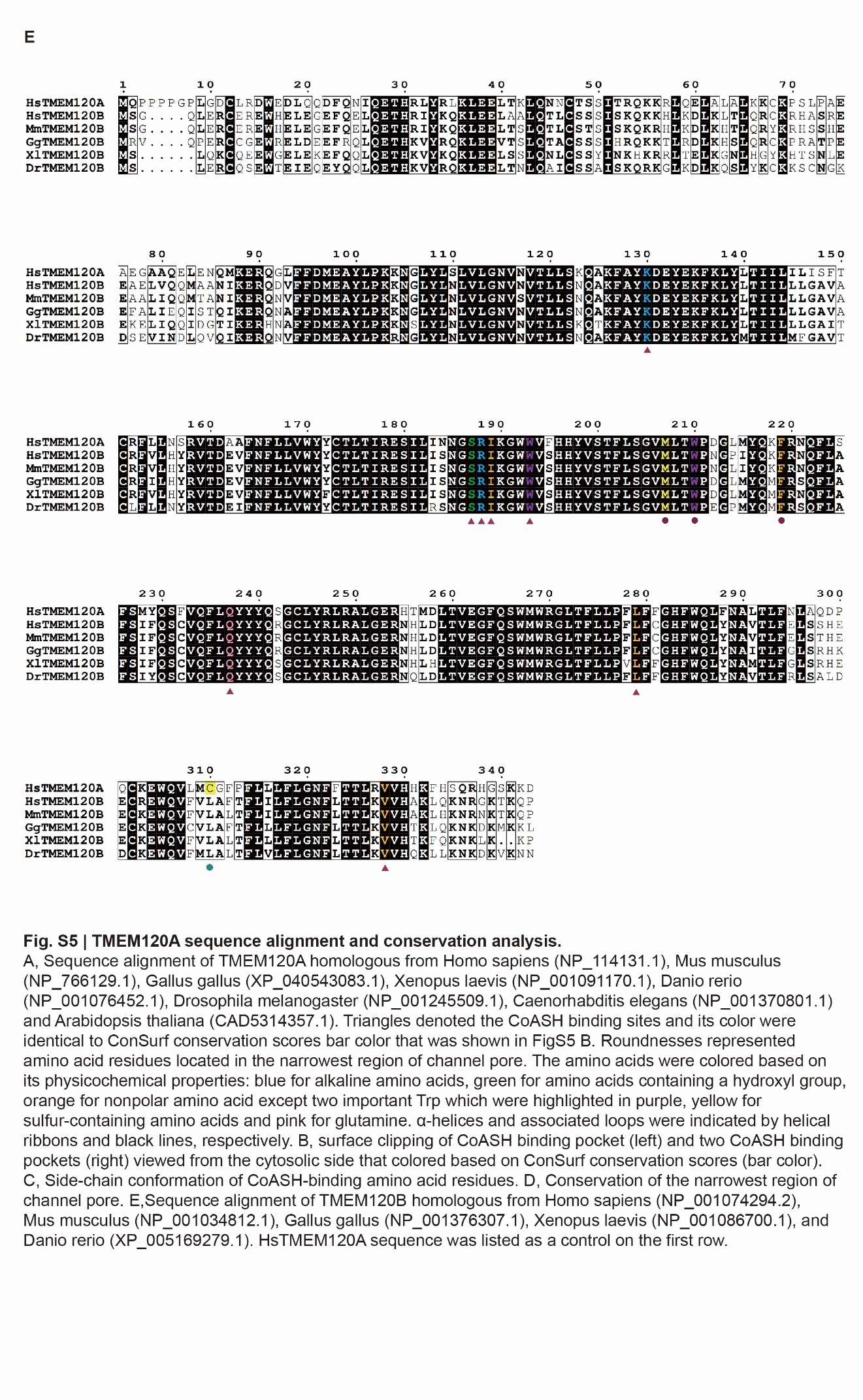

**Figure S6. Sequence alignment of *Hs*TMEM120A with various TMEM120B orthologs.** Hs, *H. sapiens* (NP_001074294.2); Mm, *M. musculus* (NP_001034812.1); Gg, *G. gallus* (NP_001376307.1); Xl, *X. laevis* (NP_001086700.1); Dr, *D. rerio* (XP_005169279.1).

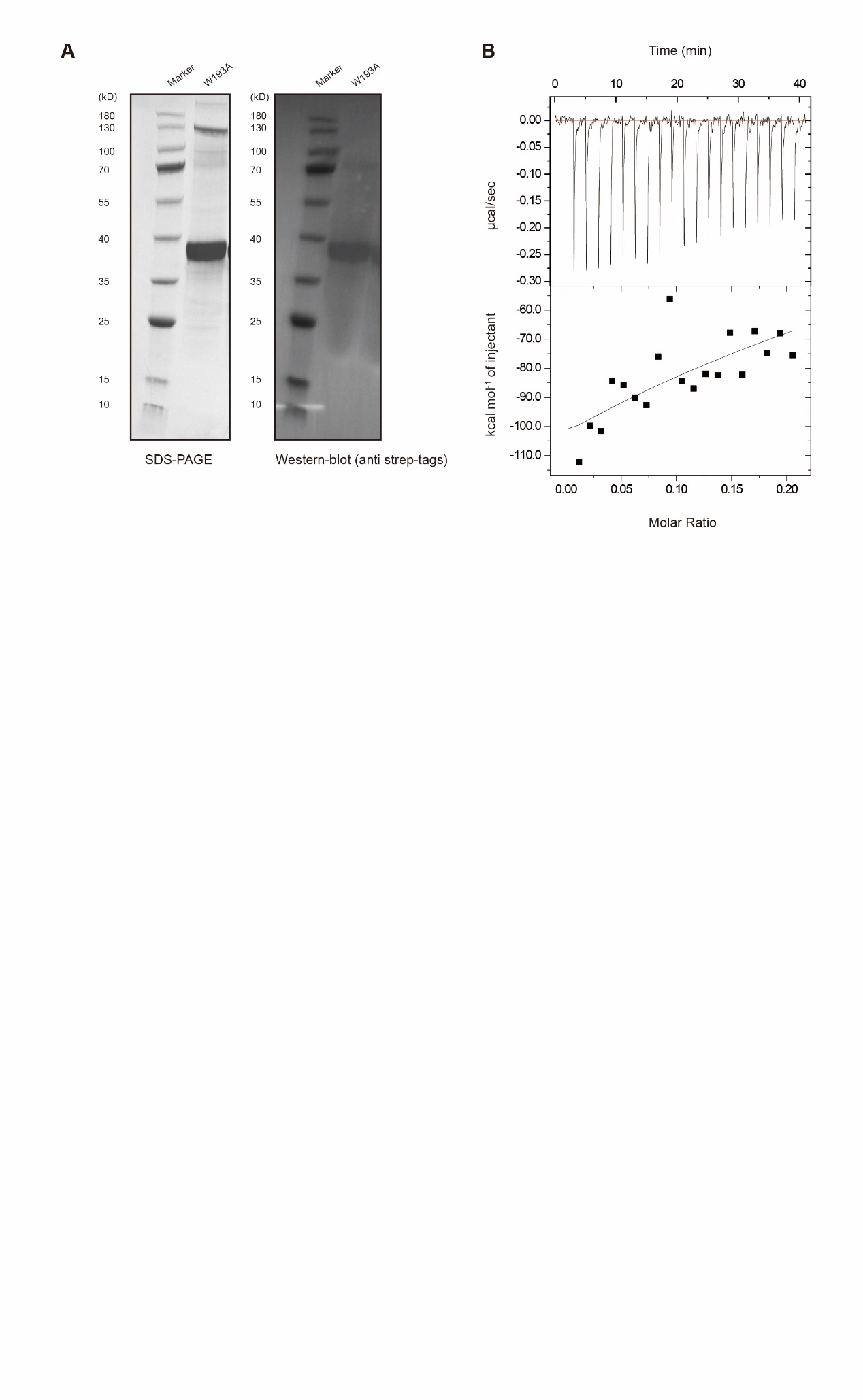

**Fig. S7. Analysis of CoASH-binding property of the W193A mutant of *Hs*TMEM120A.** **A**, SDS-PAGE and western blot analyses of the purified W193A protein. **B**, ITC analysis of W193A protein sample titrated with CoASH. The data were fit with the single-site binding isotherm model with *K*_d_ = 198.41 ± 0.81 μM.

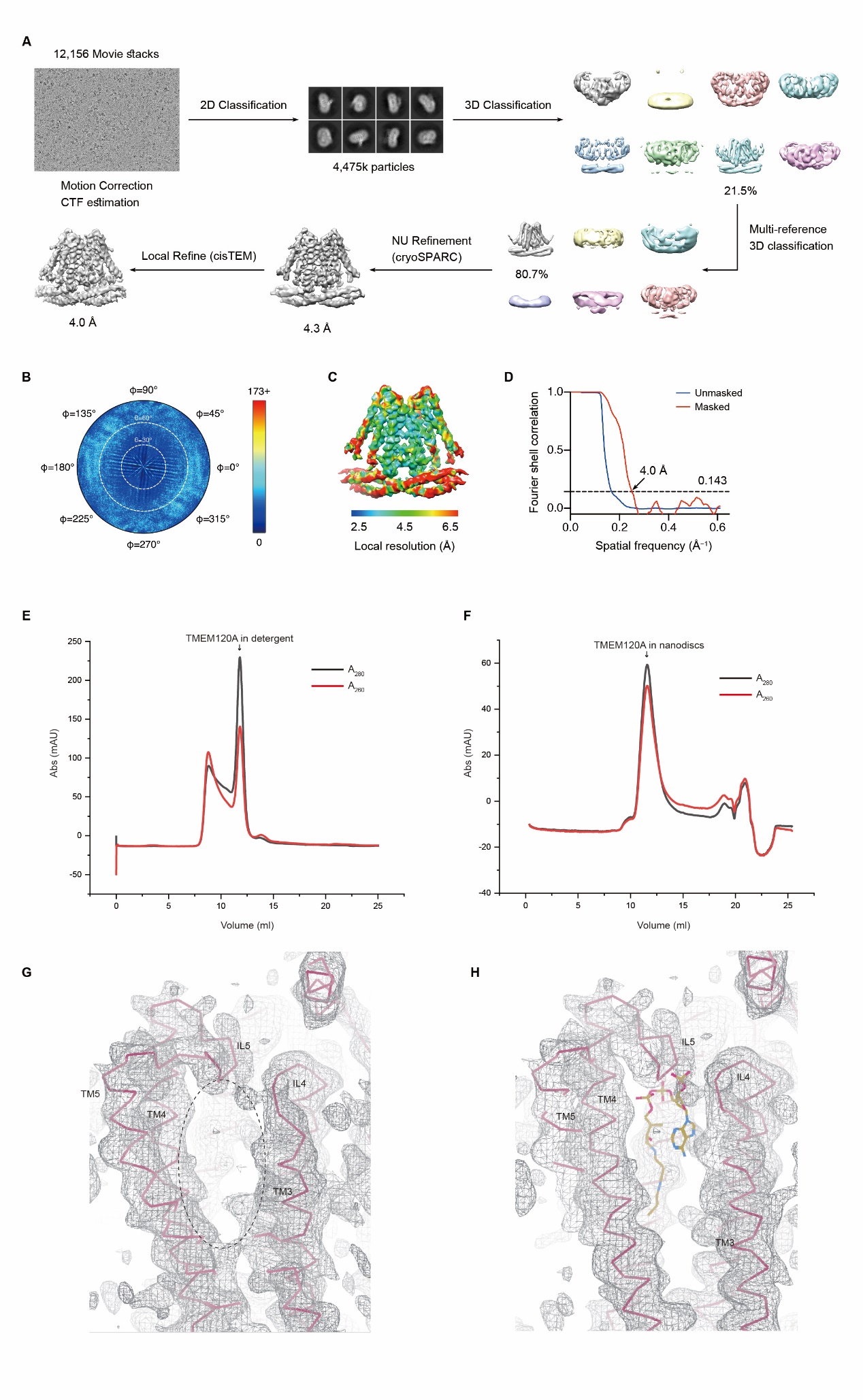

**Figure S8. Cryo-EM data collection and processing of the *Hs*TMEM120A protein purified in detergent micelle. A,** The flow chart of data collection, 2D classification, 3D classification and refinement process. A total of 12,156 movie stacks were collected, and a representative motion-corrected micrograph is shown. Particles were subjected to several rounds of 2D and 3D classifications. The final map was generated by applying local refine procedure in cisTEM. **B,** Angular distribution diagram of the particles contributing to the final reconstruction. **C,** Estimation of the local resolution in the sharpened 4.0-Å cryo-EM map of *Hs*TMEM120A protein. **D,** The FSC curves of the map with (red) or without (blue) mask applied. The resolution was estimated according to GSFSC (FSC = 0.143) criterion. **E,** Gel filtration profile of the HsTMEM120A protein eluted in detergent solution. **F,** Gel filtration profile of the HsTMEM120A reconstituted in nanodiscs. Note the apparently lower A_260_/A_280_ peak ratio in e when compared to the one in f. **G,** The local area in the CoASH-binding cavity of the HsTMEM120A structure solved in detergent. The elliptical dash ring indicates the CoASH binding site as observed in the structure of nanodisc complex. **H**, A CoASH model from the nanodisc complex structure is superposed on the map region shown in G. Note that there is no density feature accounting for the CoASH molecule in the cavity. The cryo-EM map is contoured at 0.2481 V (3.48 rmsd) level.

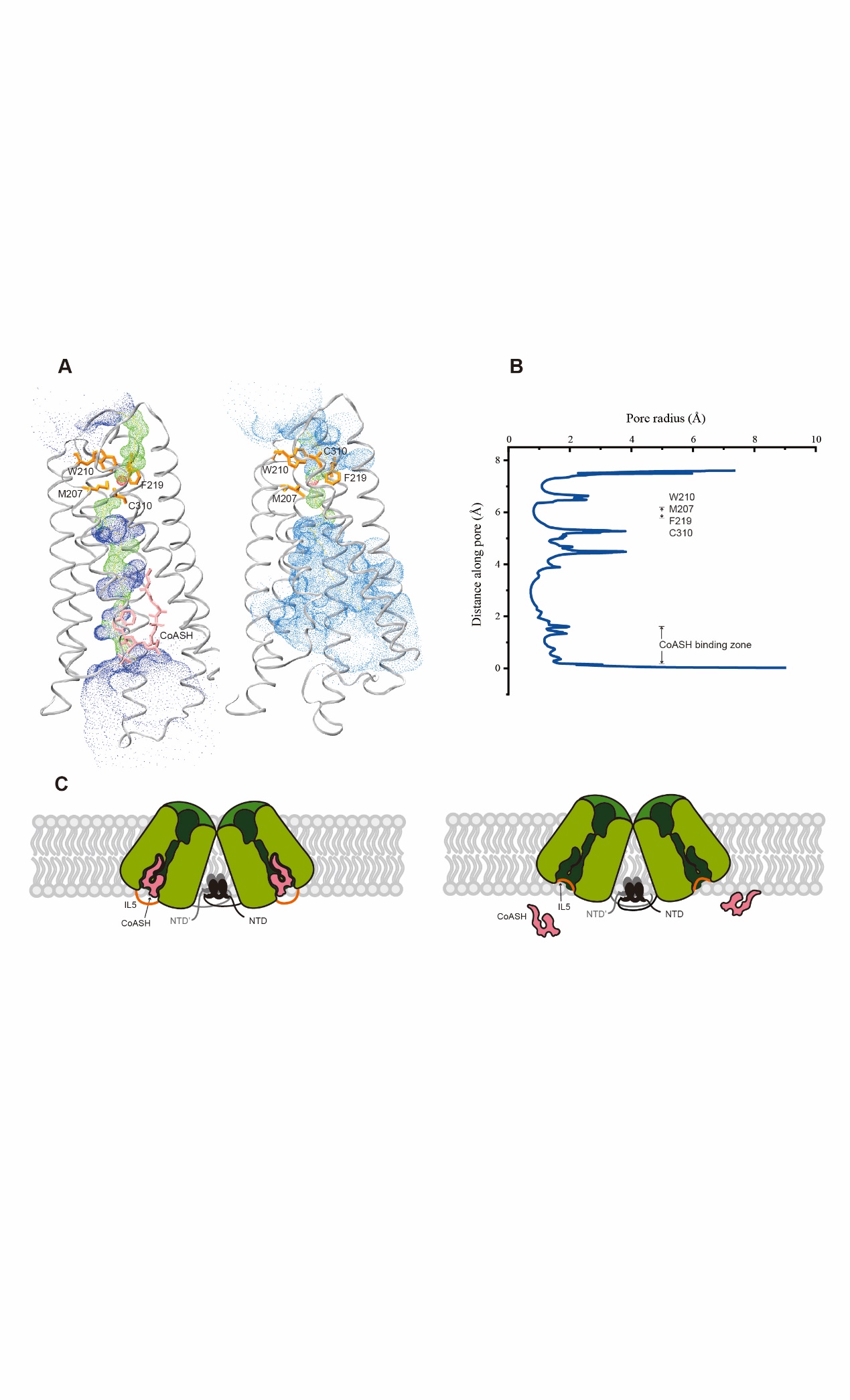

**Figure S9 | The occluded cavity of *Hs*TMEM120A with CoASH bound in comparison with the open cavity without CoASH bound. A,** Hole profiles of the CoASH-bound (left) and CoASH-free (right) *Hs*TMEM120A structures. Red, pore radius < 1.15 Å; green, 1.15 Å < pore radius < 2.30 Å; blue, pore radius > 2.30 Å. The amino acid residues surrounding the narrowest site on the extracellular side and the CoASH molecule are shown as stick models in orange and pink respectively. **B**, Distribution of the pore radius along the central axis of the *Hs*TMEM120A-CoASH complex. The constricted area around CoASH on the intracellular side and four amino acid residues on the extracellular side are labelled on the right. **C**, A mechanistic model accounting for two different functional states of TMEM120A in the cells.

**Table S1. Cryo-EM data collection and processing, refinement and validation statistics of *Hs*TMEM120A structures.**

|  | *Hs*TMEM120A in nanodiscs | *Hs*TMEM120A in detergent |
| --- | --- | --- |
|  | (EMD-31440) | (EMD-31441) |
|  | (PDB 7F3T) | (PDB 7F3U) |
| **Data collection and processing** |  |  |
| Magnification | 81,000 | 105,000 |

| Voltage (kV) | 300 | 300 |
| --- | --- | --- |
| Electron exposure (e^-^/Å) | 60 | 60 |
| Defocus range (μm) | -1.2 to -1.8 | -1.2 to -1.8 |
| Pixel size (Å) | 1.07 | 0.82 |
| Symmetry imposed | *C*2 | *C*2 |
| Initial particle images (no.) | 6,908,315 | 4,475,146 |
| Final particle images (no.) | 410,963 | 491,986 |
| Map resolution (Å) | 3.69 | 4.0 |
| FSC threshold | 0.143 | 0.143 |
| Map resolution range (Å) | 2.5-6.5 | 2.5-6.5 |
| **Refinement** |  |  |
| initial model used (PDB code) | - | - |
| Model resolution (Å) | 3.80 | 4.45 |
| FSC threshold | 0.5 | 0.5 |
| Model resolution range (Å) | - | - |
| Map sharpening *B* factor (Å^2^) | -200 | -277 |
| Model composition |  |  |
| Non-hydrogen atoms | 5632 | 4276 |
| Protein residues | 660 | 656 |
| Water | - | - |
| Ligands | 2 | - |
| *B* factors (Å^2^) |  |  |
| Protein | 52.98 | 120.11 |
| Ligand | 68.41 | - |
| R.m.s. deviations |  |  |
| Bond lengths (Å) | 0.010 | 0.006 |
| Bond angles (°) | 1.221 | 0.865 |
| Validation |  |  |
| MolProbity score | 2.05 | 2.31 |
| Clashscore | 14.73 | 20.67 |

| Poor rotamers (%) | 0.30 | 0 |
| --- | --- | --- |
| Ramachandran plot |  |  |
| Favored (%) | 94.51 | 91.72 |
| Allowed (%) | 5.18 | 8.28 |
| Disallowed (%) | 0.30 | 0 |
